# High quality genome sequence reveals the 12 pseudo-chromosomes of *Ganoderma boninense*

**DOI:** 10.1101/817510

**Authors:** Condro Utomo, Zulfikar Achmad Tanjung, Redi Aditama, Rika Fithri Nurani Buana, Antonius Dony Madu Pratomo, Reno Tryono, Tony Liwang

**Author notes:** Address correspondence to Condro Utomo. **Author’s contribution:** C.U. and T.L.: conceive the study, lead and supervise the project, Z.A.T., R.A.: perform genome assembly and analysis, R.F.N.B. and A.D.M.P.: perform fungal cultivation, R.T.: conceive the study and wrote the manuscript, All authors approved the submitted manuscript.

## Abstract

*Ganoderma boninense* is the dominant fungal pathogen in causing Basal Stem Rot (BSR) disease on oil palm. The whole genome of this fungus was sequenced using PacBio RS II platforms to gain the whole-genome shotgun libraries. These libraries were smoothed with Illumina Hiseq 4000 paired end sequencing to polish the genome assembly. Subsequently, a combination of Dovetail Chicago and HiC libraries with the HiRise assembly pipeline revealed the 12 pseudo-chromosomes of *G. boninense*. This is the first report of chromosomal-scale genome assembly for the most important fungal pathogen on oil palm.

## Introduction

The basidiomycete fungus of *Ganoderma boninense* Pat. is the causal agent of basal stem rot (BSR) disease on oil palm^1^. The disease is reported as a major economic importance on oil palm plantations in Indonesia, Malaysia, and Papua New Guinea, by reducing yield up to 50-80%^2,3^. Considering its economic impact, a draft genome sequence was assembled to complement the lack of genomic information of this fungus using Pacific Biosciences (PacBio) and Illumina platforms for the Indonesian strain (G3)^4^. However, the chromosome level assembly for this fungus remains uncovered.

Current sequencing technologies and assembly algorithms change the genome sequencing landscape. Chromosomal assembly can be done through single-molecule sequencing and chromatin conformation capture (3C) data analysis^5–7^. PacBio and Illumina sequencing lay the fundamental layer of sequence scaffolds contiguity and resolve repetitive DNA regions^8,9^. The Chicago technique improves scaffolding of de novo PacBio and Illumina sequencing through chromatin reconstructing usage as a substrate to obtain proximity ligation libraries *in vitro*^10^. Additionally, the HiC which is an adaptation of 3C approach, is used to detect chromatin interaction in the nucleus through *in vivo* formaldehyde-crosslink and physically form 3D architecture of the genome^11^. The HiRise algorithm assembly pipeline in both Chicago and HiC analysis anchors, orders, and orients the assembly sequences within each chromosome. In this study, we reported the first chromosome genome assembly of *G. boninense* using PacBio long-reads, Illumina short-reads, Chicago sequencing and HiC technology.

## Methods

### Sample collection

A *G. boninense* G3 strain was isolated from an oil palm tree with severe symptoms of BSR disease in North Sumatera Province, Indonesia. Freshly pure revived mycelia were grown in 100 ml yeast malt broth in dark at 28°C for 14 days. Mycelia were harvested on a layer of Whatman paper no. 1 and air dried for 15 minutes. Half of the sample was proceeded for genomic DNA isolation using GenElute plant genomic DNA miniprep kit (Sigma-Aldrich Co., St. Louis, MO, USA) according to the manufacturer’s instructions for PacBio and Illumina platforms sequencing. Another part of the sample, mycelia were freeze-dried and shipped for Chicago and HiC platforms sequencing.

### PacBio/Illumina sequencing and de novo assembly

Single-molecule sequencing was performed using PacBio RS II with the latest P6-C4 chemistry systems and Illumina HiSeq 4000 system according to each manufacturer’s instruction. The PacBio reads was assembled using WTDBG2 with default parameters^12^. Subsequently, two rounds of Racon was used for consensus calling for this assembly^13^. At the final assembly step, two rounds of PacBio-Racon assemblies were polished by basecall correction through Pilon software with default parameters^14^. Pilon uses Illumina reads to perform base corrections and derive an accurate consensus sequence.

### Chicago library preparation and sequencing

A Chicago library was prepared as described previously^10^. Briefly, ∼500ng of high molecular weight of genomic DNA (gDNA) (mean fragment length = 59) was reconstituted into chromatin *in vitro* and fixed with formaldehyde. Fixed chromatin was digested with DpnII, the 5’ overhangs filled in with biotinylated nucleotides, and then free blunt ends were ligated. After ligation, crosslinks were reversed and the DNA purified from protein. Purified DNA was treated to remove biotin that was not internal to ligated fragments. The DNA was then sheared to ∼350 bp mean fragment size and sequencing libraries were generated using NEBNext Ultra enzymes and Illumina-compatible adapters. Biotin-containing fragments were isolated using streptavidin beads before PCR enrichment of each library. The libraries were sequenced on an Illumina HiSeq X to produce 165 million 2×150 bp paired end reads, which provided 1,887.11 × physical coverage of the genome (1-100 kb pairs).

### Dovetail HiC library preparation and sequencing

A Dovetail HiC library was prepared in a similar manner as described previously^15^. Briefly, for each library, chromatin was fixed in place with formaldehyde in the nucleus and then extracted. Fixed chromatin was digested with DpnII, the 5’ overhangs filled in with biotinylated nucleotides, and then free blunt ends were ligated. After ligation, crosslinks were reversed and the DNA purified from protein. Purified DNA was treated to remove biotin that was not internal to ligated fragments. The DNA was then sheared to ∼350 bp mean fragment size and sequencing libraries were generated using NEBNext Ultra enzymes and Illumina-compatible adapters. Biotin-containing fragments were isolated using streptavidin beads before PCR enrichment of each library. The libraries were sequenced on an Illumina HiSeq X to produce 194 million 2×150 bp paired end reads, which provided 41,650.61 × physical coverage of the genome (10-10,000 kb pairs).

### Scaffolding the assembly with HiRise

The input *de novo* assembly, shotgun reads, Chicago library reads, and Dovetail HiC library reads were used as input data for HiRise, a software pipeline designed specifically for using proximity ligation data to scaffold genome assemblies^10^. An iterative analysis was conducted. First, Shotgun and Chicago library sequences were aligned to the draft input assembly using a modified SNAP read mapper (http://snap.cs.berkeley.edu). The separations of Chicago read pairs mapped within draft scaffolds were analyzed by HiRise to produce a likelihood model for genomic distance between read pairs, and the model was used to identify and break putative misjoins, to score prospective joins, and make joins above a threshold. After aligning and scaffolding Chicago data, Dovetail HiC library sequences were aligned and scaffolded following the same method. After scaffolding, shotgun sequences were used to close gaps between contigs.

## Results and Discussion

The single-molecule genome sequencing of the G3 strain was performed using PacBio RS II platforms combined with Illumina Hiseq 4000 paired end sequencing technology to obtain the whole-genome shotgun libraries. For PacBio RSII platform, a 20 kb library was built and sequenced. Simultaneously, paired-end reads were generated by sequencing of initial libraries of 300 bp using Illumina HiSeq4000 system.

The 3C analysis was conducted using sequencing-based Chicago and HiC assemblies. Using Chicago approach, Library 1 produced 165 million reads of 2×150 bp and provided 1,887.11 × physical coverage of the genome (1-100 kb pairs). In HiC approach, Library 1 produced 194 million read pairs of 2×150 bp and provided 41,650.61 × physical coverage of the genome (10-10,000 kb pairs).

Furthermore, the libraries from PacBio and Illumina were assembled using WTDBG2, Racon, and Pilon to produce 55.82 Mb with 592 scaffolds and N50 value of 0.357 Mb. Both Chicago and HiC libraries were assembled with HiRise pipeline. Using PacBio and Illumina as an input data, the Chicago HiRise assembly resulted in L90/N90 of 75 scaffolds. This result was used as an input for HiC HiRise assembly and resulted the L90/N90 output into 12 scaffolds (**Table 1**). The contiguity comparison graph showed the significance improved of assembly (**Figure 1**).

**Table 1.**
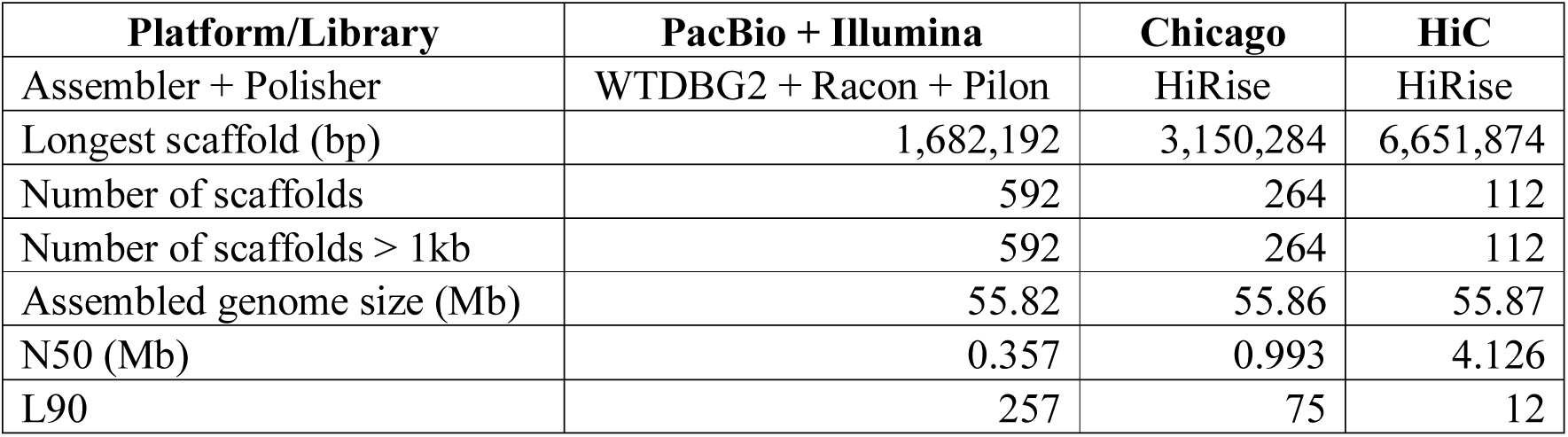
Comparison of de novo and proximity-guided genome assembly.

**Figure 1.**
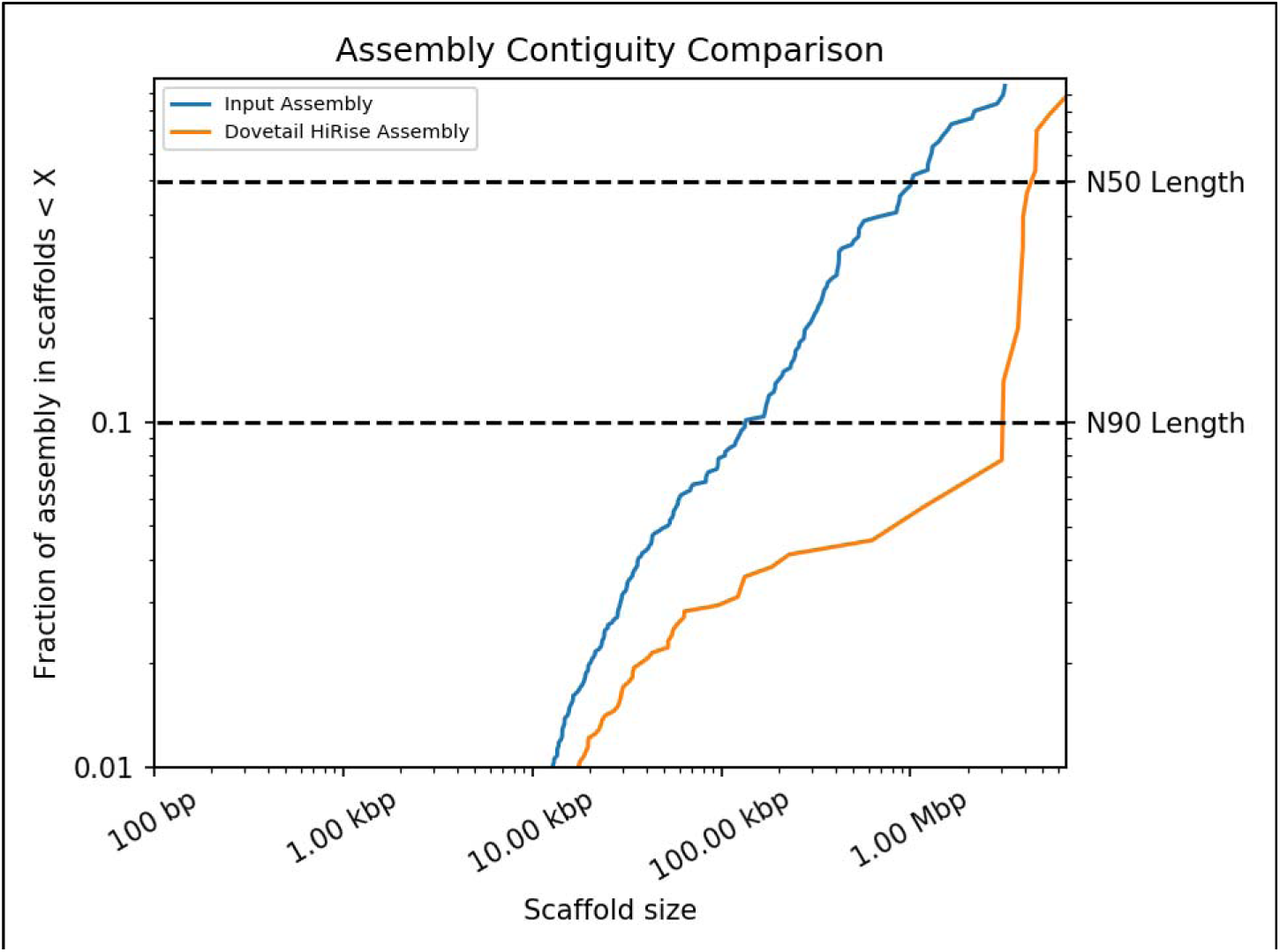
A comparison of the contiguity of the input assembly and the final HiRise scaffolds. Each curve shows the fraction of the total length of the assembly present in scaffolds of a given length or smaller. The fraction of the assembly is indicated on the Y-axis and the scaffold length in basepairs is given on the X-axis. The two dashed lines mark the N50 and N90 lengths of each assembly. Scaffolds less than 1 kb are excluded.

The improvement of scaffolds assembly occurred from the role of the usage of biotin-labeled nucleotide in HiC which enables selective purification of chimeric DNA ligation junctions^11^. HiC data provides spanning even whole chromosomes to improve scaffold contiguity of assemblies chromosome-length scaffolds for genomes^16^.

In addition, Dovetail Genomics’ HiRise pipeline generated a HiC linkage density histogram plot (**Figure 2**). The plot compares the positions of read pair sequences (the pair of end sequences from each and every sequenced DNA fragment obtained by chromatin cross-linking) versus the positions of each individual DNA sequence within the genome assembly^10,17^. The alignment produces a diagonal of lines from lower left to upper right in the plot that represent each of the 12 pseudo-chromosomes. Dots (sequences) within boxes at the last column are probably un-scaffold DNA sequence.

**Figure 2.**
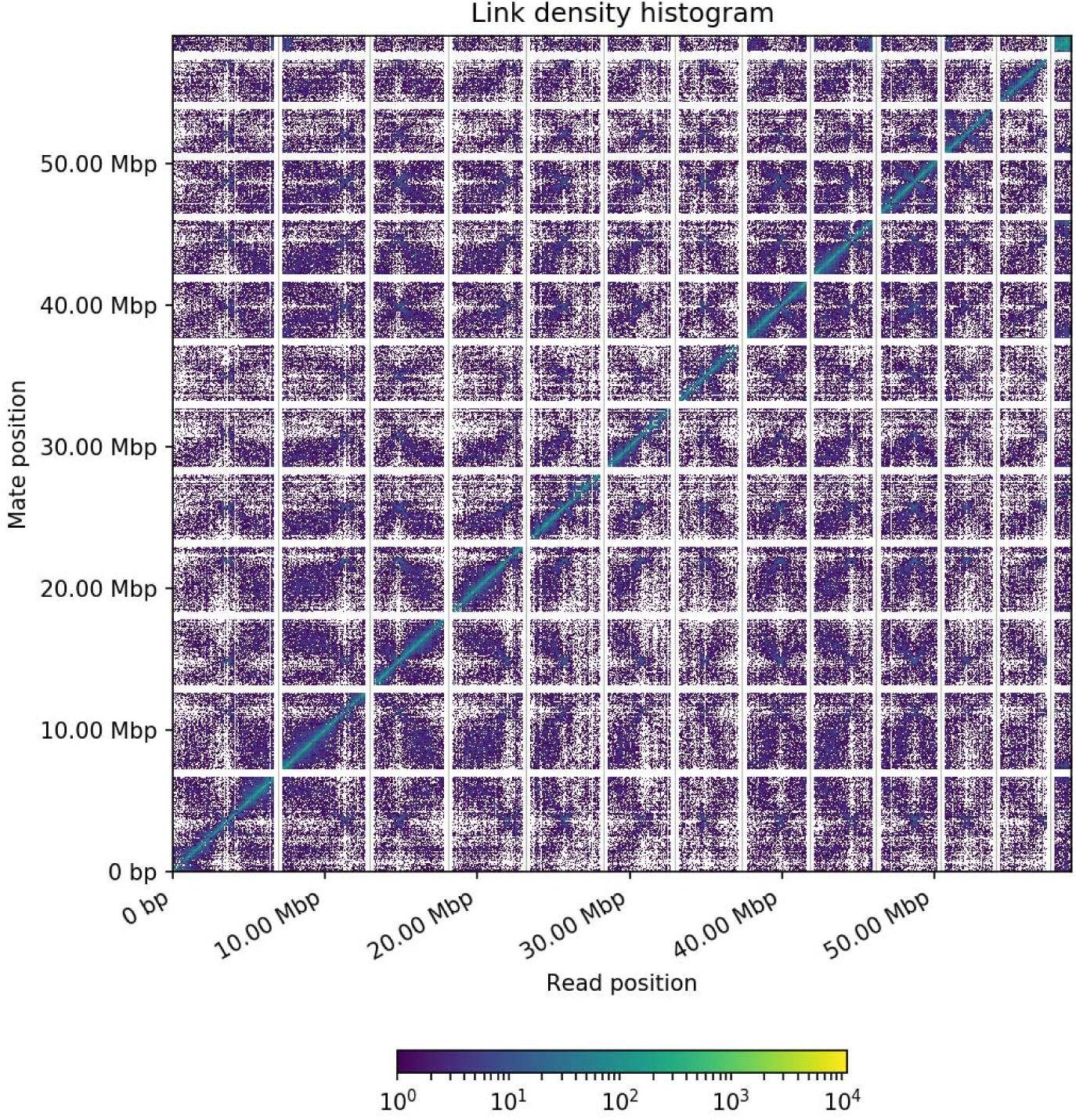
Dovetail Genomic’s HiC linkage density histogram. In this figure, the x and y axes give the mapping positions of the first and second read in the read pair respectively, grouped into bins. The color of each square gives the number of read pairs within that bin. White vertical and black horizontal lines have been added to show the borders between scaffolds. Scaffolds less than 1 Mb are excluded.

Overall, the single-molecule sequencing combined with chromatin cross-linking deep sequencing resulted in 12 scaffolds. The density histogram plot is able to guide the anchoring of scaffolds to pseudo-chromosomes^18,19^. It is thus corresponding to the pseudo-chromosome number in *G. boninense*. This number is resembling two other close-related *Ganoderma* species that the chromosomal assembly has been generated earlier i.e. *G. lucidum* (13 chromosomes) and *G. sinense* (12 chromosomes)^20,21^. In term of gene number, *G. boninense* is the highest with 21,074 coding sequences followed with *G. lucidum* with 16,113, and *G. sinense* with 15,688 genes. Gene density of each pseudo-chromosome in *G. boninense* is scattered evenly in all pseudo-chromosomes (**Figure 3**). This study showed that the final HiC assembly provides a robust, fast, and valid data for generating de novo assemblies with chromosome-length scaffolds. However, these assemblies still contain 504 gaps (Table 2).

**Table 2.** Assembly statistics for HiC libraries and HiRise assembly of *G. boninense*.

| Characteristics | Number |
| --- | --- |
| Number of pseudo-chromosome | 12 |
| Number of scaffold | 112 |
| Longest scaffold (Mb) | 6.5 |
| N50 scaffold (Mb) | 304.34 |
| L50 (scaffolds) | 6 |
| N90 (Mb) | 3.032 |
| L90 (scaffolds) | 12 |
| Length of genome assembly (Mb) | 55.87 |
| GC contents (%) | 56.2 |
| Number of protein-coding genes | 21,074 |
| GC content of protein-coding genes ( % ) | 59.2 |
| Average gene length (bp) | 1.265 |
| Average number of exons per gene | 7 |
| Average exon size (bp) | 178 |
| Average intron size (bp) | 97 |
| Average size of intergenic regions (bp) | 1.250 |
| Number of gaps | 504 |
| Non-gap bases (bp) | 55.821.405 |

**Figure 3.**
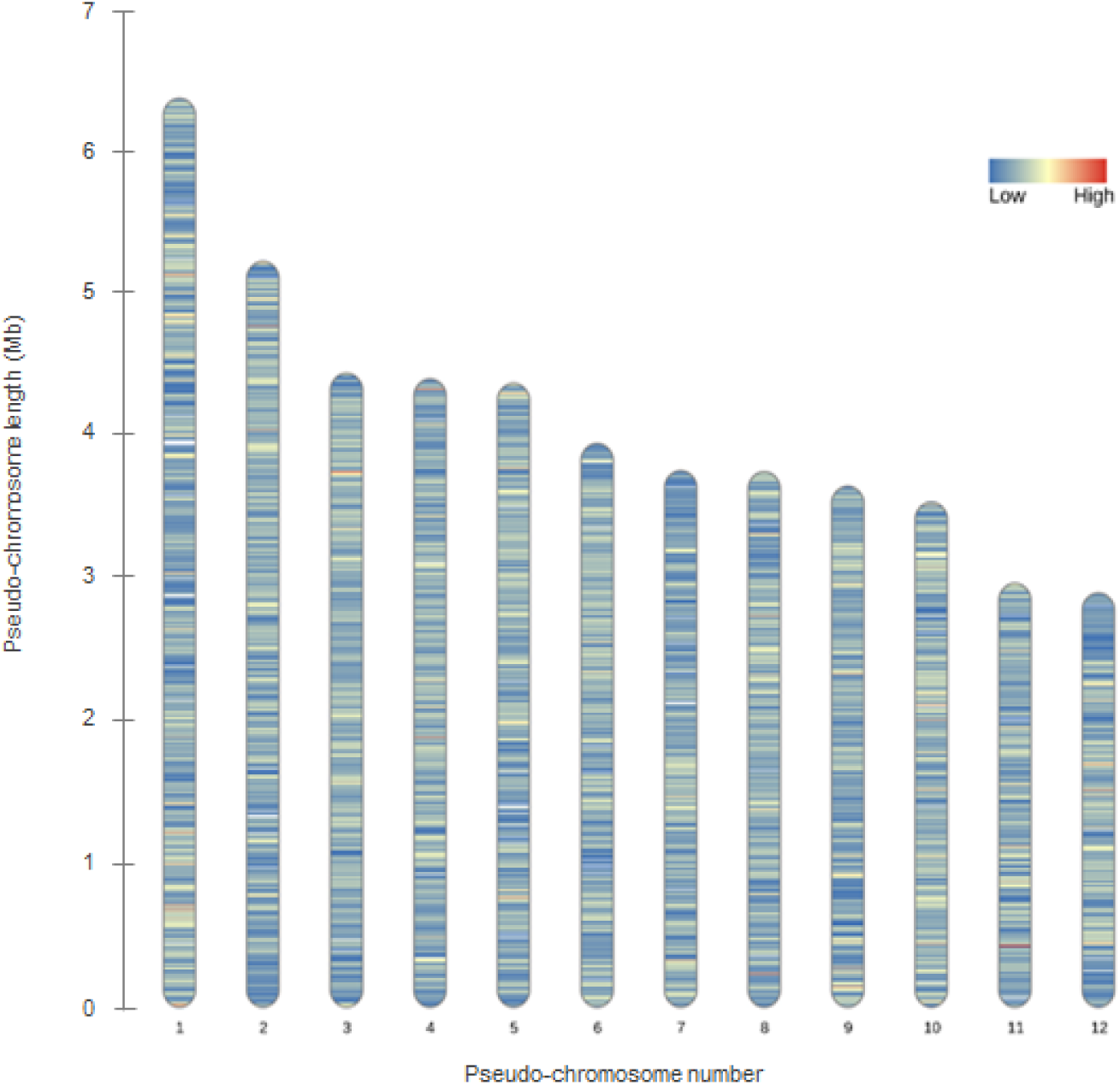
Gene density distribution in all pseudo-chromosomes of *G. boninense*. This graph excludes 100 scaffolds with total of 4.3 Mb length that unmapped which size ranges between 1.1 Mb and 2.6 Kb.

## Conclusion

This study demonstrates a chromosomal-scale genome assembly of *G. boninense* through combination of single-molecule sequencing (PacBio long reads and Illumina short reads) and 3C data analysis (Chicago and HiC with HiRise pipeline). These combined technologies harbored 55.87 MB length of genome assembly within 12 pseudo-chromosomes and an additional 4.3 Mb un-scaffold sequences.

## Acknowledgement

This work was supported by the management of PT SMART Tbk. We thank Roberdi, Marcelinus Rocky Hatorangan, and Victor Aprilyanto for proof-read this manuscript, and Sanju Rianintika and Hani Feorani for their technical supports.

